# Myelination deficits in the auditory brainstem of a mouse model of Fragile X Syndrome, a possible mechanism for auditory phenotypes

**DOI:** 10.1101/2021.09.08.459530

**Authors:** Alexandra Lucas, Shani Poleg, Achim Klug, Elizabeth McCullagh

## Abstract

Auditory symptoms are one of the most frequent sensory issues described in people with Fragile X Syndrome (FXS), the most common genetic form of intellectual disability. However, the mechanisms that lead to these symptoms are under explored. In this study, we examined whether alterations in myelination in auditory brainstem circuitry contribute to auditory impairments in FXS mice. Specifically, we studied myelinated fibers that terminate in the Calyx of Held, which encode temporally precise sound arrival time, and are some of the most heavily myelinated axons in the brain. We measured anatomical myelination characteristics using coherent anti-stokes Raman spectroscopy (CARS) and electron microscopy (EM) in a FXS mouse model in the medial nucleus of the trapezoid body (MNTB) where the Calyx of Held synapses. We measured number of mature oligodendrocytes (OL) and oligodendrocyte precursor cells (OPCs) to determine if changes in myelination were due to changes in the number of myelinating or immature glial cells. The two microscopy techniques (EM and CARS) showed a decrease in fiber diameter in FXS mice. Additionally, EM results indicated reductions in myelin thickness and axon diameter, and an increase in g-ratio, a measure of structural and functional myelination. Lastly, we showed an increase in both OL and OPCs in MNTB sections of FXS mice suggesting that the myelination phenotype is not due to an overall decrease in number of myelinating OLs. This is the first study to show that a potential myelination mechanism can explain alterations seen in FXS at the level of the auditory brainstem.

## 2 Introduction

Fragile X Syndrome (FXS) is the most common monogenic form of autism spectrum disorder (ASD) and occurs in 1:4000-1:8000 people in the U.S. (Hagerman and Hagerman, 2008). FXS has been commonly used as a model for studying ASD, especially because there are several commercially available animal models such as mouse and rat (The Dutch-Belgian Fragile X Consorthium et al., 1994; Till et al., 2015; Tian et al., 2017). There have been many proposed mechanisms for how loss of Fragile X Mental Retardation Protein (FMRP), the protein encoded by the gene *Fmr1*, leads to FXS and ASD phenotypes (Bear et al., 2004; Gantois et al., 2017; Hagerman et al., 2009; Osterweil et al., 2013; Rajaratnam et al., 2017, among others). However, drug therapies in animal models do not always “rescue” human phenotypes when extended to the clinic (Dahlhaus, 2018). Recent work has shown that myelination may underly some the phenotypes common in ASD and FXS (Pacey et al., 2013; Phan et al., 2020). Auditory phenotypes, such as auditory hypersensitivity, are often conserved between mouse and human FXS, and it is possible that myelination deficits are also conserved (McCullagh et al., 2020b; Phan et al., 2020). There is reduced or delayed myelination in FXS in many brain areas (Pacey et al., 2013; Phan et al., 2020) and one of the targets of FMRP is myelin basic protein (Jeon et al., 2017), suggesting that deficits seen in FXS may be caused at least in part by alterations to myelination. Additionally, research has shown FMRP expression in <u>mature</u> oligodendrocytes (OLs), potentially explaining how loss of FMRP impacts myelination (Wang et al., 2004; Giampetruzzi et al., 2013). The mechanisms by which binaural hearing is impaired in people with FXS and ASD are unknown, however there is evidence the auditory brainstem is involved (Kulesza and Mangunay, 2008; Kulesza Jr. et al., 2011; Wang et al., 2014; Rotschafer et al., 2015; Garcia-Pino et al., 2017; McCullagh et al., 2017, 2020a; Zorio et al., 2017; Curry et al., 2018; El-Hassar et al., 2019; Lu, 2019).

Previous work has shown that FXS impacts the auditory brainstem and sound localization processing in a myriad of ways, from protein dysfunction to behavioral changes (reviewed in (McCullagh et al., 2020b). FMRP expression is high in the brainstem, suggesting an important role in brainstem function (Wang et al., 2014; Zorio et al., 2017). Alterations to proteins which directly interact with FMRP can critically disrupt the maintenance of neuronal activity patterns in the brainstem (Brown et al., 2010; Strumbos et al., 2010; El-Hassar et al., 2019). In addition, previous anatomical and physiological work has shown that excitation/inhibition balance within the auditory brainstem circuit is altered in FXS (Rotschafer et al., 2015; Garcia-Pino et al., 2017; McCullagh et al., 2017; Rotschafer and Cramer, 2017; Curry et al., 2018; Lu, 2019). This circuit relies on precise timing of excitatory and inhibitory inputs for computation of sound location information (reviewed in Grothe et al., 2010). Behavioral studies have shown subtle, but persistent, auditory specific behavioral phenotypes in FXS mice that likely originate in altered auditory brainstem processing (McCullagh et al., 2020a) however the exact mechanism that underlies these alterations is unknown.

The ability to localize sound is dependent on a very precise comparison of sound input from the two ears, either through interaural timing differences (ITD) or interaural sound intensity/level differences (IID). Processing and interpretation of ITD and IID relies on extremely fast and temporally precise synaptic transmission, leading to some of the most heavily myelinated axons in the brain (Ford et al., 2015). In the auditory brainstem, the medial nucleus of the trapezoid body (MNTB) afferent axons stand out due to their large diameters, which are about ∼ 3 μm in healthy hearing animals (Ford et al., 2015). The large diameter is due to a high demand on speed and temporal precision by this synapse (Kim et al., 2013b). Any alteration in this pathway that changes the precision and timing of this circuit will lead to substantial difficulties in the ability to separate competing auditory streams and localize sound, as occurs in ASD. Interestingly, increased changes to latency for both behavioral and physiological measures seem to be one of the more repeatable phenotypes in FXS mice (Kim et al., 2013a; McCullagh et al., 2020a, but see Rotschafer et al., 2015; El-Hassar et al., 2019). One way in which changes to myelination in the auditory brainstem would likely manifest is through changes in the speed of synaptic transmission, and therefore latency of auditory brainstem timing.

Several studies have shown that myelination deficiencies contribute to ASD phenotypes in several brain areas. This suggests that an area of the brain most critically dependent on myelination for properly encoding sound location information, such as the auditory brainstem, should be especially impacted (Pacey et al., 2013; Phan et al., 2020). We use a combination of microscopy techniques (coherent anti-stokes Raman spectroscopy (CARS) and electron microscopy (EM)) to examine fine myelin microstructure in Fmr1 KO mice compared to wildtype controls (B6). Additionally, we measured the number of mature and precursor oligodendrocytes (OLs and OPCs) as a potential mechanism through which myelin microstructure is altered in FXS mice. We hypothesize that there will be reduced myelin (thickness and diameter) and concomitant changes in number of OLs and OPCs in FXS mice. The goal of this study is to provide a possible mechanism through which auditory phenotypes might arise in FXS.

## 3 Materials and Methods

All experiments complied with all applicable laws, National Institutes of Health guidelines, and were approved by the University of Colorado Anschutz Institutional Animal Care and Use Committee.

### 3.1 Animals

All experiments were conducted in either C57BL/6J (stock #000664, B6) background or hemizygous male and homozygous *Fmr1* knock-out strain (B6.129P2-*Fmr1*^*tm1Cgr*^/J stock #003025, Fmr1 KO) obtained from The Jackson Laboratory (Bar Harbor, ME USA)(The Dutch-Belgian Fragile X Consorthium et al., 1994). Animals from both sexes were used in the experiments for both B6 and Fmr1 KO mice. There were no significant differences in any of the measures based on sex, so data for both sexes are combined for analysis (p-values shown are the main effect of sex; CARS p = 0.7454, OL count p = 0.9529, OPC count p = 0.2103, EM data was one animal of each sex so differences do not apply here). Exact number of animals used are listed in the figure legends for each experiment but ranged from 4-7 for each genotype and were between P72-P167 (postnatal day).

### 3.2 Tissue preparation

Mice were overdosed with pentobarbital (120 mg/kg body weight) and transcardially perfused with phosphate-buffered saline (PBS; 137 mM NaCl, 2.7 mM KCl, 1.76 mM KH2PO4, 10 mM Na2HPO4 Sigma-Aldrich, St. Louis, MO, USA) followed by 4% paraformaldehyde (PFA). After perfusion, the animals were decapitated, and the brains removed from the skull. Brains were kept overnight in 4% PFA before transferring to PBS. Brainstems were embedded in 4% agarose (in PBS) and sliced coronally using a Vibratome (Leica VT1000s, Nussloch, Germany) at either 200 µm thickness for myelination analysis with CARS or 70 µm thickness for labeling oligodendrocytes.

For EM imaging, mice were perfused with PBS followed by a solution containing 2.5% glutaraldehyde, 2% paraformaldehyde, and 0.1 M cacodylate buffer at pH 7.4. The brains were stored in the same solution for 24 hours and sectioned at 500 µm using the same protocol as above. The sections were immersed in glutaraldehyde solution for a minimum of 24 hours at 4° C. The tissue was then processed by rinsing in 100 mM cacodylate buffer followed by immersion in 1% osmium and 1.5% potassium ferrocyanide for 15 min. Next, the tissue was rinsed five times in the cacodylate buffer and immersed again in 1% osmium for 1.5 hrs. After rinsing five times for 2 min each in cacodylate buffer and two times briefly in water, e*n bloc* staining with 2% uranyl acetate was done for at least 1 hour at 4 ° C, followed by three rinses in water. The tissue was transferred to graded ethanols (50, 70, 90, and 100%) for 15 minutes each and then finally to propylene oxide at room temperature, after which it was embedded in LX112 and cured for 48 hrs at 60 ° C in an oven. Ultra-thin parasagittal sections (55 nm) were cut on a Reichert Ultracut S (Leica Microsystems, Wetzlar, Germany) from a small trapezoid positioned over the tissue and were picked up on Formvar-coated slot grids (EMS, Hatfield, PA, USA).

### 3.3 Immunohistochemistry

For staining oligodendrocytes, six to eight free-floating sections from each brain were submerged in L.A.B. solution (Liberate Antibody Binding Solution, Polysciences, Inc, Warrington, PA, USA, Cat No. 24310) for ten min to help expose epitopes related to labeling mature and precursor oligodendrocytes. Next, the slices were washed two times in PBS and blocked in a solution containing 0.3% Triton-X (blocking solution), 5% normal goat serum (NGS) and PBS for one hr on a laboratory shaker. After blocking, slices were stained with primary antibodies rabbit anti-Aspartoacylase (GeneTex, Irvine, CA, USA; 1:1000) and mouse Sox-10 (A-2) (Santa Cruz, Dallas, TX, USA; sc-365692, 1:500) diluted in blocking solution with 1% NGS and incubated overnight (Table 1). Slices were then washed three times (10 min each wash) in PBS and incubated in secondary antibodies (Table 2) diluted in blocking solution with 1% NGS for two hrs. After three additional washes in PBS (5 min each wash), Nissl (Neurotrace 425/435 Blue-Fluorescent Nissl Stain, Invitrogen, Carlsbad, CA USA, 1:100) in antibody media (AB media: 0.1 M phosphate buffer (PB: 50 mM KH2PO4, 150 mM Na2HPO4), 150 mM NaCL, 3 mM Triton-X, 1% bovine serum albumin (BSA)) was applied for thirty min. Stained slices were then briefly washed in PBS and slide-mounted with Fluoromount-G (Southern Biotech, Cat.-No.: 0100-01, Birmingham, AL, USA). Slides were stored at 4°C. Slices used for myelination analysis with CARS (four to five slices per brain) were stained with Nissl as described above, then stored free-floating at 4°C until imaged. All antibody labeling was performed at room temperature.

**Table 1:**
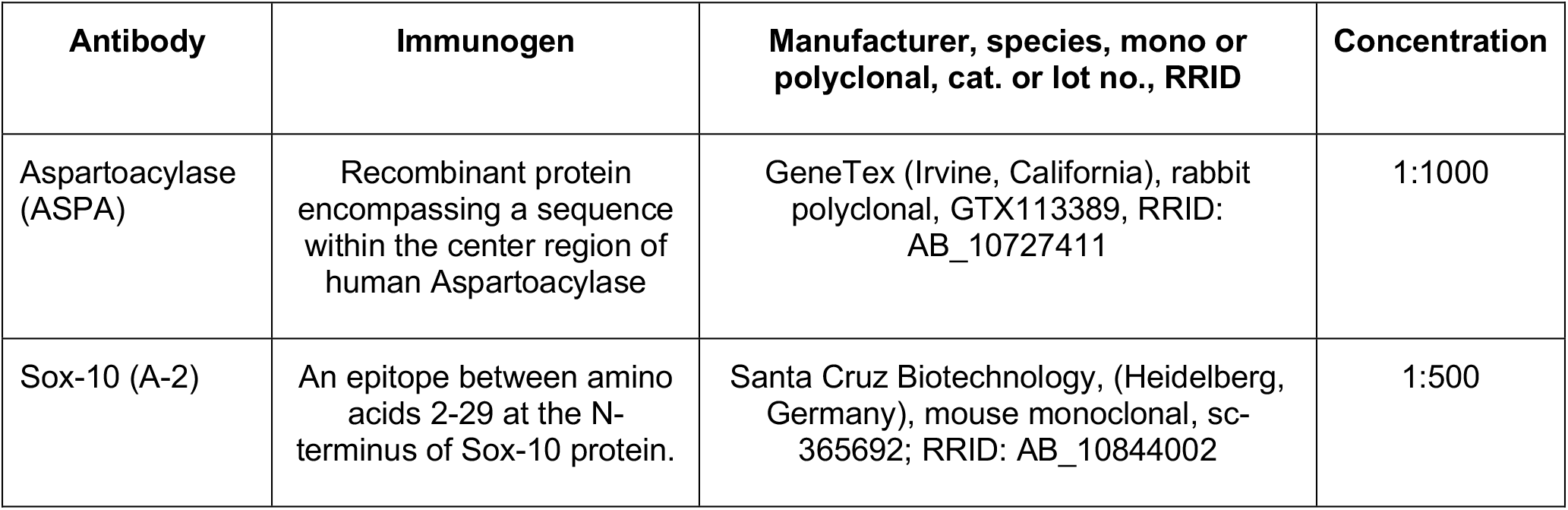
Primary antibodies used in immunofluorescence.

**Table 2:** Secondary antibodies used in immunofluorescence.

| Catalog No., RRID | Manufacturer | Reactivity | Conjugate | Concentration |
| --- | --- | --- | --- | --- |
| A-21428, AB_2535849 | Invitrogen | Goat anti-Rabbit IgG (H+L) | Alexa Fluor 555 | 1:500 |
| A-21235, AB_2535804 | Invitrogen | Goat anti-Mouse IgG (H+L) | Alexa Fluor 647 | 1:500 |

### 3.4 Antibody Characterization

The primary antibody for mature oligodendrocytes (Aspartoacylase (ASPA), 1:1000, GeneTex, Irvine, CA, USA; GTX113389; RRID: AB_10727411, Table 1) is a rabbit polyclonal antibody specific to mature oligodendrocytes (OLs). The ASPA antibody is specific to a recombinant protein encompassing a sequence within the center region of the human aspartoacylase. Aspartoacylase is responsible for the conversion of N-acetyl_L-aspartic acid (NAA) to aspartate and acetate. Hydrolysis by aspartoacylase is thought to help maintain white matter (Bitto et al., 2007). The ASPA antibody has been shown to label mature oligodendrocytes in mouse cerebral cortex and has been validated by protein overexpression and western blot analysis (Larson et al., 2018; Orthmann-Murphy et al., 2020). Additional western blot analyses with various whole cell extracts also showed that ASPA antibody detects aspartoacylase protein. The primary antibody Sox-10 (Sox-10 (A-2), 1:500, Santa Cruz Biotechnology, Heidelberg, Germany; sc-365692; RRID: AB_10844002) was used to label the entire oligodendrocyte lineage (oligodendrocyte precursor cells (OPCs) and OLs). Sox-10 is a mouse monoclonal antibody specific to an epitope mapping between amino acids 2-29 at the N-terminus of the human Sox-10 gene. Sox-10 has been shown to specifically label the oligodendrocyte lineage in the CNS and in mouse brains (Zuo et al., 2018; Barak et al., 2019; Imamura et al., 2020). Both primary antibodies were visualized using two fluorescent-conjugated secondary antibodies which are listed in Table 2.

### 3.5 Imaging

Brainstem slides for immunofluorescence were imaged using an Olympus FV1000 confocal microscope (Olympus, Tokyo, Japan) with lasers for 488, 543 and 635 nm imaging. Once MNTB was identified by the distinctive trapezoidal morphology and cell size, z-stacks were taken using a 20x objective (UPLSAPO20X, NA 0.75) so that the entire nucleus could be visualized and quantified. The nucleus was separated into medial, central, and lateral regions following the protocol in Weatherstone et al., 2016. Briefly, the MNTB was digitally extracted using FIJI software, and the tonotopic axis was estimated by drawing the longest possible dorsomedial-to-ventrolateral line through the nucleus (Weatherstone et al., 2016).This line was measured using FIJI and divided into thirds wherein perpendicular lines were drawn. These lines delineated the lateral, central and medial regions of MNTB. For CARS microscopy, brainstem sections were imaged using an Olympus FV-1000 (Olympus Corp., Tokyo, Japan) fitted with non-descanned detectors in both the forward and epi CARS directions. Sections were placed in a culture dish with coverslip (for inverted microscopy) and custom weight to keep tissue near coverslip. Z-stacks were taken with a 60X, 1.2NA infrared corrected water objective which served for collection of the CARS signal in the epi direction for medial and lateral MNTB. Signal in the forward direction was collected through an Olympus 0.55NA condenser. The APE picoEmerald laser set up contained an NKT fiber laser which provided the 80MHz clock, and an OPO (Optical Parametric Oscillator) laser with a tunable range of 770-990nm. Water absorption peak was at 1388cm^-1^ which implies a 901.1 nm pump/probe beam with the fixed Stokes beam at 1031nm and a CARS signal at 801.6nm. To excite the CH2 stretch at 2845cm^-1^ the APE picoEmerald lasers required a pump/probe beam set to 797.2 nm which resulted in a CARS signal at 649.8 nm, thus laser settings were always set to 797.2 nm prior to imaging. Pulse duration for the two lasers was approximately 2 ps and the polarization was linear and horizontal. The synchronization was primarily based on the OPO cavity length with the pulse pumped by the frequency doubled output from the NKT laser at 515.5 nm. The tunable pump/probe beam was used for the TPEF, which was separated by a dichroic from the CARS signal. The NDD epi-direction unit contained two PMTS of the same type.

Sections for EM were first imaged on a compound microscope, and MNTB was identified by the characteristic cell size and shape, as well as the location of the cells in the parasagittal slices. Once MNTB location was determined, slices were imaged on a FEI Tecnai G2 transmission electron microscope (Hillsboro, OR, USA) with an AMT digital camera (Woburn, MA, USA).

### 3.6 Cell counting

Quantification of OL’s and entire OL lineage (oligodendrocyte precursor cells (OPCs) and OLs) was performed using the optical fractionator approach and FIJI software (Schindelin et al., 2012). In a pilot study to determine the appropriate stereological parameters, five brains were sliced at 50 µm using a freezing microtome (SM2010R Leica, Nussloch, Germany) and z-stacks from 5 sections per brain (every other section) were taken with a 20x objective the same as above. Using FIJI each stack was brightened, the background subtracted, the image scaled and the MNTB was digitally extracted. Separate grid plugins were used to create the dissector and count frames for each image stack. To account for tissue shrinkage, the section thickness was measured at multiple points within MNTB using the Olympus FV1000 microscope and FV10-ASW microscopy software. Measurements were taken by recording the first z-level where tissue features came into focus, and then recording the last z-level where tissue features were in focus. The difference between the recorded levels for each measured area of MNTB was averaged and represented the section thickness for stereological calculations. Stacks were set to contain 20 slices regardless of the section thickness and the step size was kept constant within, but not between brains. Cells were counted by moving through the stack and using a cell counter plugin to mark the top of the cell when it first came into focus within the probe unless it contacted the exclusion lines of the count frame (Schmitz and Hof, 2005).

The pilot data indicated that selecting four 70 µm sections (every other section starting from a randomly selected slice) to give a section subfraction (SSF) of 0.280 was appropriate for calculating the total number of cells in a single MNTB. In addition, counts would use a probe area of (1250 µm × 1250 µm) and dissector (5000 µm × 5000 µm) giving an area subfraction (ASF) of 0.25. It should be noted that two brains had fewer than 4 slices (3.5 and 3 slices respectively) which were suitable for quantification. OL analyses also included three brains from the pilot study that had adequate immunohistochemical staining for quantification. In these cases, the SSF was adjusted, or a fifth slice was counted to achieve an SSF that was closest to 0.280 (0.20-0.25). The height subfraction (HSF) was calculated separately for each brain using the mean dissector height (OL mean = 19.5 µm - 34.4 µm; OPC mean = 21.06 µm - 34.4 µm) and the mean section thickness (OL mean = 26.75 µm - 45.75 µm; OPC mean = 27.75 µm – 45.75 µm). Using these parameters, OL’s and OPC’s were counted and recorded in excel. OPC’s were identified as cells that were labeled with Sox 10, but not co-labeled with ASPA (Figure 1). Calculation of the total cell count for each complete MNTB nucleus was done by multiplying the counted cells (∑Q^-^) with the reciprocal sampling fractions such that:

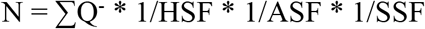

**Figure 1.**
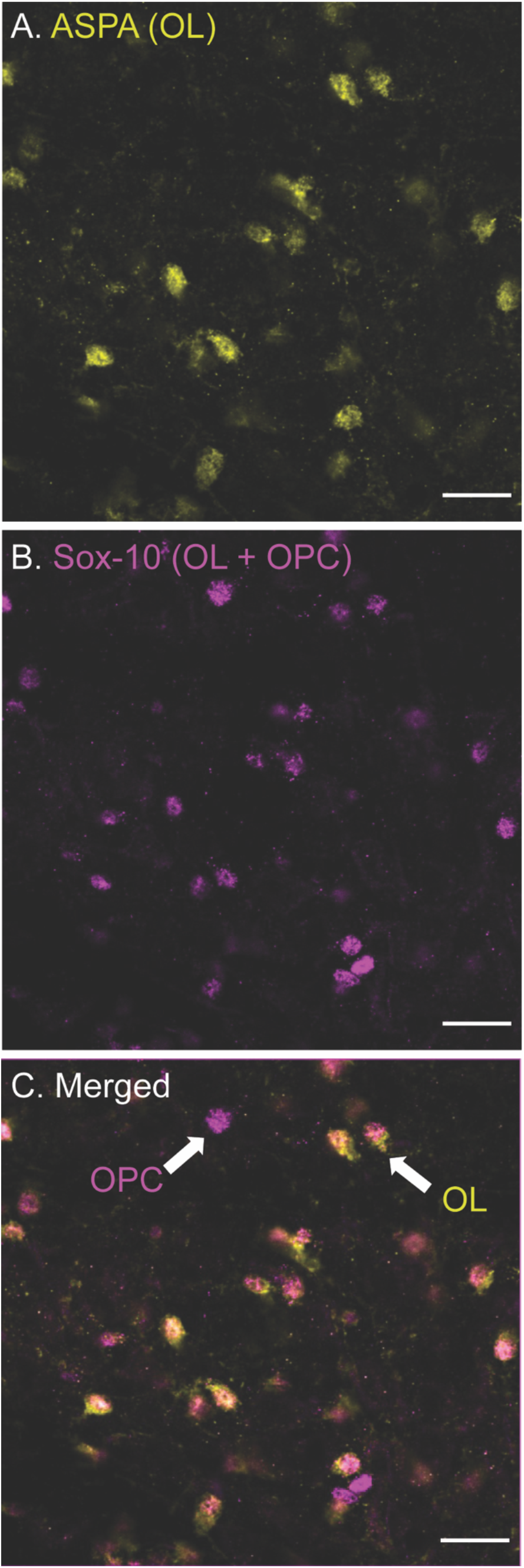
Differentiation of oligodendrocyte (OLs, ASPA) and oligodendrocyte precursor cells (OPCs, Sox-10 no ASPA). ASPA, a stain specific for mature OLs is shown in yellow (panel A). These yellow cells were used to calculate the number of mature OLs in each image. Sox-10 is a marker specific for the entire OL lineage (OPCs and OLs) and is labeled in magenta (B). To determine the number of OPCs in each image, cells that were labeled with Sox-10 but not ASPA (as indicated by the arrow labeled OPC) were counted (C).

Error in the stereological approach was calculated using the Gundersen-Jensen coefficient of error (CE) estimator and smoothness class of m=1 (Gundersen et al., 1999). CE measures error as a function of the biological variance (noise) and the sampling variance from systemic uniformly random sampling. The coefficient of variance (CV) within and between brains was high (CV>1) due to natural variations in the population of glial cells throughout MNTB. CE for OL and OPC estimates was greater than 10% (Mean CE = 0.51), but the sampling parameters chosen showed CE was negligible in terms of overall variation (CE^2^ /CV^2^ < 0.003)(Gundersen et al., 1999). Thus, the stereological parameters were determined to be sufficient for quantification estimates.

### 3.7 Myelination analysis

Analysis of myelination morphology was done using FIJI software. Myelin diameter was measured from CARS images by using the line tool to measure the fiber diameter (Figure 2A) for 20 axons per image. The resolution was not sufficient in CARS images to quantify thickness or g-ratio of axons. Tissue shrinkage was not corrected for in this analysis. In addition, diameters of 240 axons were also measured in the same way in EM images, however myelin thickness was measured directly, and g-ratios were calculated from fiber and axon diameters (Figure 2B)(Ford et al., 2015). G-ratios were calculated from EM images using inner and outer fiber diameter ratio (g-ratio = inner fiber diameter / outer fiber diameter). EM data is representative of one animal from each genotype and was performed primarily to confirm CARS diameter changes.

**Figure 2.**
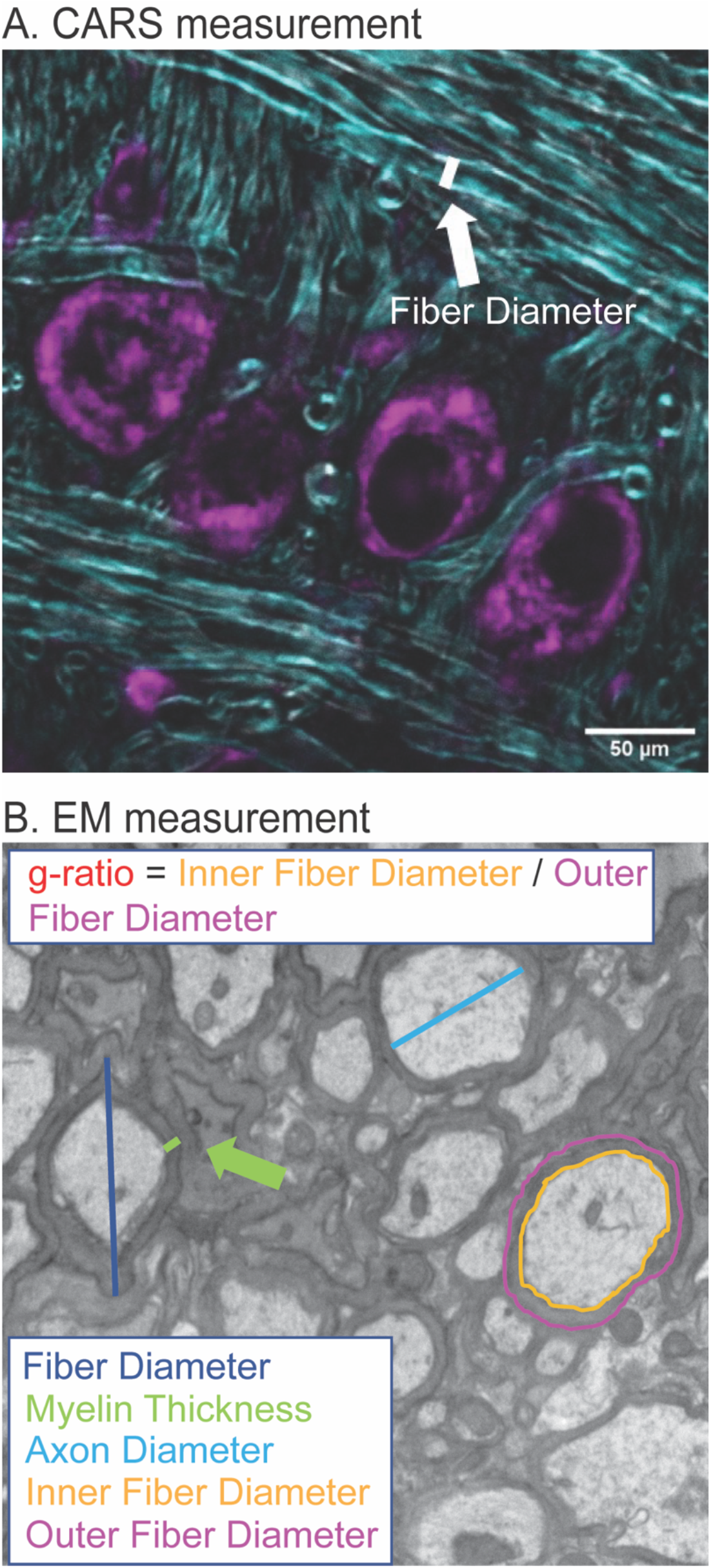
Quantification of myelination morphology in CARS and EM images. CARS images were quantified using the FIJI line segment tool. A line, as indicated by white line (A), was drawn and the measure function used to measure the fiber diameter for 20 axons per image. For EM images, increased resolution allowed for more detailed analysis including fiber diameter (dark blue), axon diameter (cyan), myelin thickness (green), inner (orange) and outer (purple) fiber diameter (B). Inner and outer fiber diameter were then used to calculate g-ratio, which is the inner fiber diameter/outer fiber diameter.

### 3.8 Statistical analyses

Figures were generated in R (R Core Team, 2013) using ggplot2 (Wickham, 2016). Data for all experiments were analyzed using linear mixed-effects models (lme4; (Bates et al., 2014)). To account for repeated measures and variability with animals, Animal was considered a random effect with fixed effects of genotype and location (medial, center, lateral, and total when appropriate). Additional analyses used fixed effects of age or sex to determine if these were contributing factors in the differences shown. It was expected that there may be differences between different regions of the MNTB (medial, center, lateral, total) therefore a priori, it was determined that estimated marginal means (emmeans; (Lenth, 2019)) would be used for pairwise comparisons between region and genotype. To control for multiple comparisons, emmeans implements a Tukey method for these contrasts. Tables XXX show the summary statistics for each experiment. Asterisks are used to indicate statistical significance between the two genotypes, as follows: \**p* < 0.05, \*\**p* < 0.01, and \*\*\**p* < 0.001. Figures were prepared for publication using Photoshop and Illustrator (Adobe, San Jose, CA, USA).

## 4 Results

We used several (EM and CARS) microscopy techniques to examine the myelin microstructure of putative globular bushy cell axons in the auditory brainstem that innervate the MNTB and form the Calyx of Held in Fmr1 KO mice compared to wildtype controls (B6). We also counted the number of both mature and precursor oligodendrocytes in the same region to show if structural changes are due to changes in number or type of oligodendrocytes which myelinate these axons.

### 4.1 Myelin diameter as measured by CARS

We first examined the diameter of axons in the medial and lateral MNTB using CARS microscopy (Figure 3). CARS microscopy for myelination imaging was used over electron microscopy (EM) due to its speed and compatibility with immunofluorescence (Wang et al., 2005). We measured axon fiber diameters in both medial and lateral MNTB because previous work has shown the potential for differences in myelin based on tonotopic location (medial or lateral) (Ford et al., 2015; Stange-Marten et al., 2017). Representative images for both location (medial or lateral columns) for B6 and Fmr1 KO (rows) are shown in Figure 3A-D. We found that there was a significant decrease in fiber diameter in Fmr1 KO compared to B6 for both medial and lateral MNTB (Figure 3E). Not surprisingly when data were summed between medial and lateral locations, there was still a significant decrease in fiber diameter in Fmr1 KO mice.

**Figure 3.**
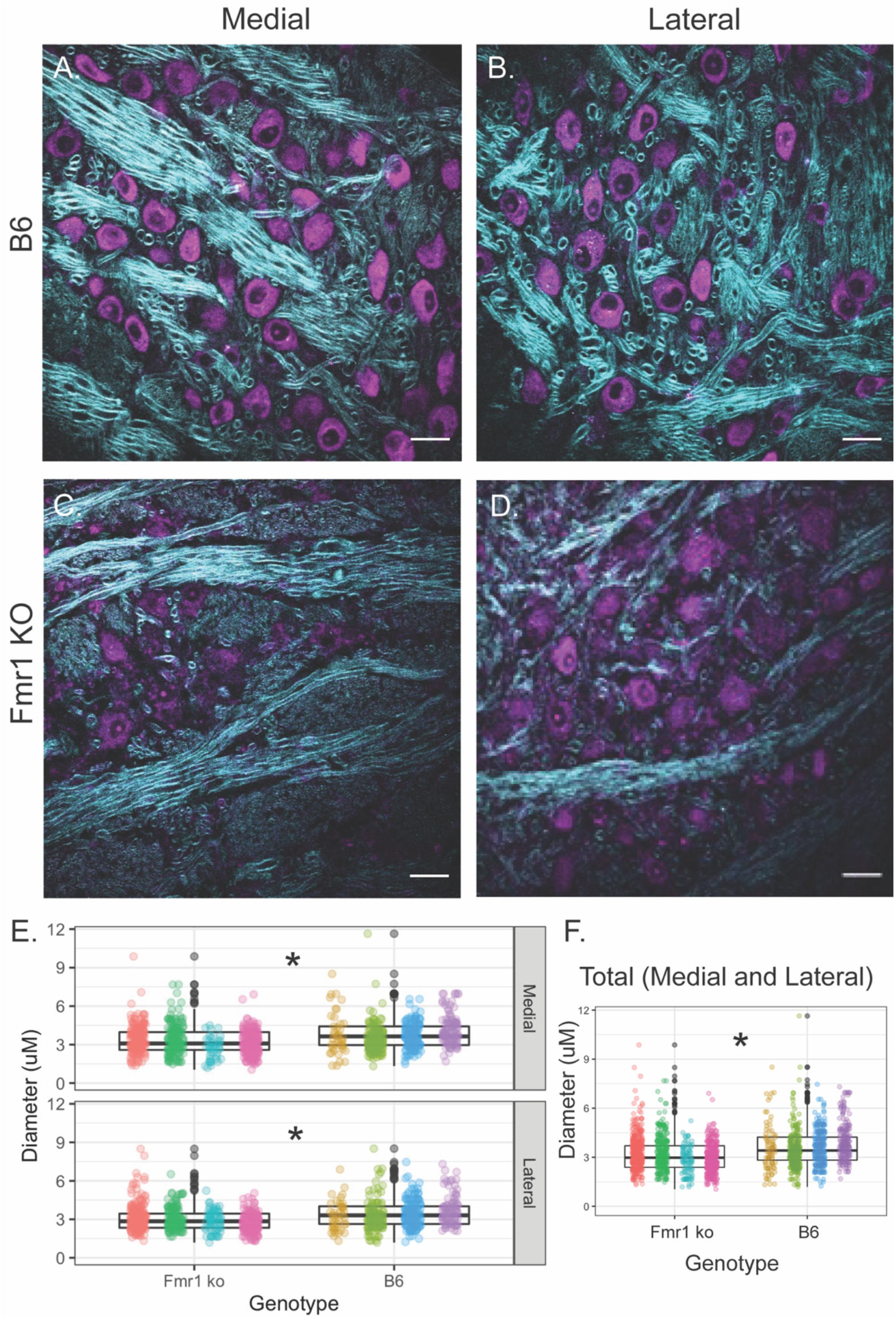
Fiber diameter size is reduced in Fmr1 KO mice compared to B6 using CARS. Representative images for medial and lateral (columns) MNTB in B6 and Fmr1 KO mice (rows)(A – D). Scale bar is 20 µm. Box plots show data from each individual animal (color spread) and population data represented by the box for Fmr1 KO (left) and B6 (right) for either medial (top) or lateral (bottom) MNTB (E). N for each genotype was 4 animals with 20 measurements per section and 4-6 sections per animal. F shows the data independent of medial or lateral location (total) for each animal across genotype. * = *p* < 0.05.

### 4.2 CARS Myelin characteristics confirmed by EM

To confirm that the CARS technology was sensitive enough to determine differences between fiber diameters in Fmr1 KO and B6 mice, we also performed EM controls on one animal of each genotype (across multiple sections with multiple measurements, Figure 4). The advantage of EM, while costly and time consuming, is that it allows for higher resolution and thus more detailed description of changes to myelination than CARS microscopy techniques. Since alterations in fiber diameter seen in CARS data was not dependent on medial or lateral localization, and it is difficult to determine location with parasagittal EM sections, these data are not separated by location. Representative EM images for B6 and Fmr1 KO are shown in Figure 4A and 4B respectively. Consistent with CARS analysis, EM data showed smaller fiber diameters in Fmr1 KO animals compared to B6 (Figure 4C). Additionally, axon diameter and myelin thickness are also decreased in Fmr1 KO compared to B6 (Figure 4D, E). Lastly, g-ratio, an often-used index of optimal functional and structural myelination, was increased in Fmr1 KO animals compared to B6 (Figure 4F, Chomiak and Hu, 2009). These results are consistent with the relatively large axon diameters in the Fmr1 KO mice compared to fiber diameters (a measure for g-ratio) despite thinner myelin and smaller axons than B6 mice.

**Figure 4.**
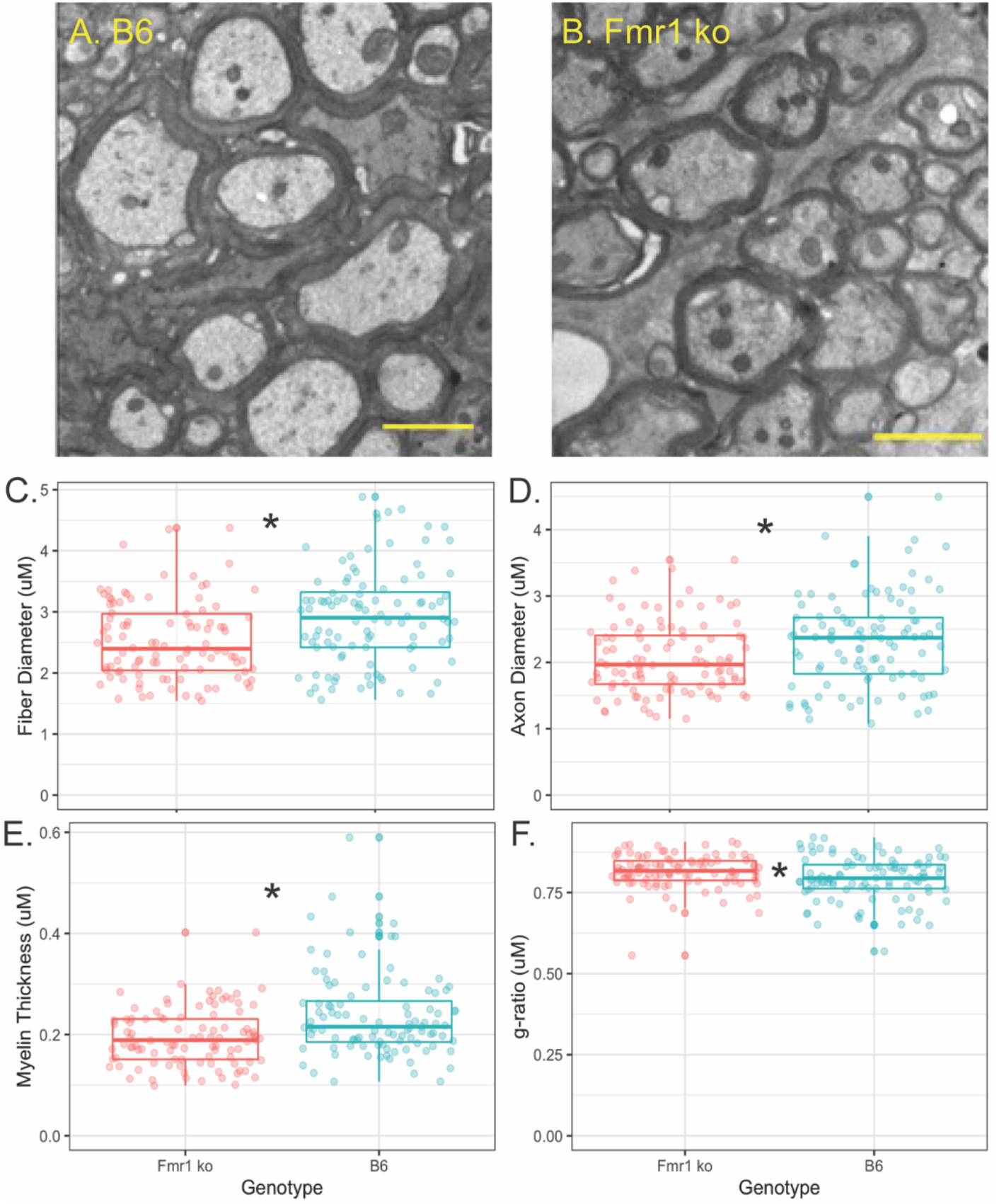
Myelin characteristics are also altered in Fmr1 KO compared to B6 mice measured by EM. Representative images for EM in B6 (A) and Fmr1 KO (B) mice. Scale bar is 2 µm. Box plots show data from each individual measurement for Fmr1 KO (left) and B6 (right) mice. Measured parameters include fiber diameter (C), axon diameter (D), myelin thickness (E), and g-ratio (F). N is one animal per genotype with multiple sections and multiple measurements / section represented by individual puncta. * = *p* < 0.05.

### 4.3 Number of mature oligodendrocytes

Oligodendrocytes (OLs) are the glial cells responsible for myelinating axons in the central nervous system. FMRP has been shown to be expressed in OLs, potentially providing a mechanism through which loss of FMRP can lead to changes in myelination (Wang et al., 2004; Giampetruzzi et al., 2013). For OL counts, the MNTB was divided into three sections, medial, central, and lateral. Figure 5A and B show representative images of the entire MNTB stained with ASPA (a marker for mature OLs) and sox-10 (a marker for the entire OL lineage) for B6 and Fmr1 KO mice respectively. As shown by arrowheads, OLs are represented by staining with ASPA (which also coincides with sox-10 staining). Cells that are only stained for sox-10 (magenta) and not ASPA (yellow) are quantified as OPCs (see below and Figure 1, 6). Like CARS data, there was no difference in OL number based on specific tonotopy of the MNTB (medial, center, or lateral) however, when data was combined for all regions, there was a significant increase in OL number in Fmr1 KO mice compared to B6 (Figure 5C top panel). Using the sampling Q count technique, to estimate number of cells within the entire MNTB similar results were found to raw counts. Specifically, there was no difference in medial, center, or lateral estimates, but total combined regions showed a highly significant (*p* <0.0001) increase in OLs in Fmr1 KO animals compared to B6. In addition to changes in number of mature OLs, another possible mechanism underlying alterations to myelination could be incomplete or impaired maturation of OLs.

**Figure 5.**
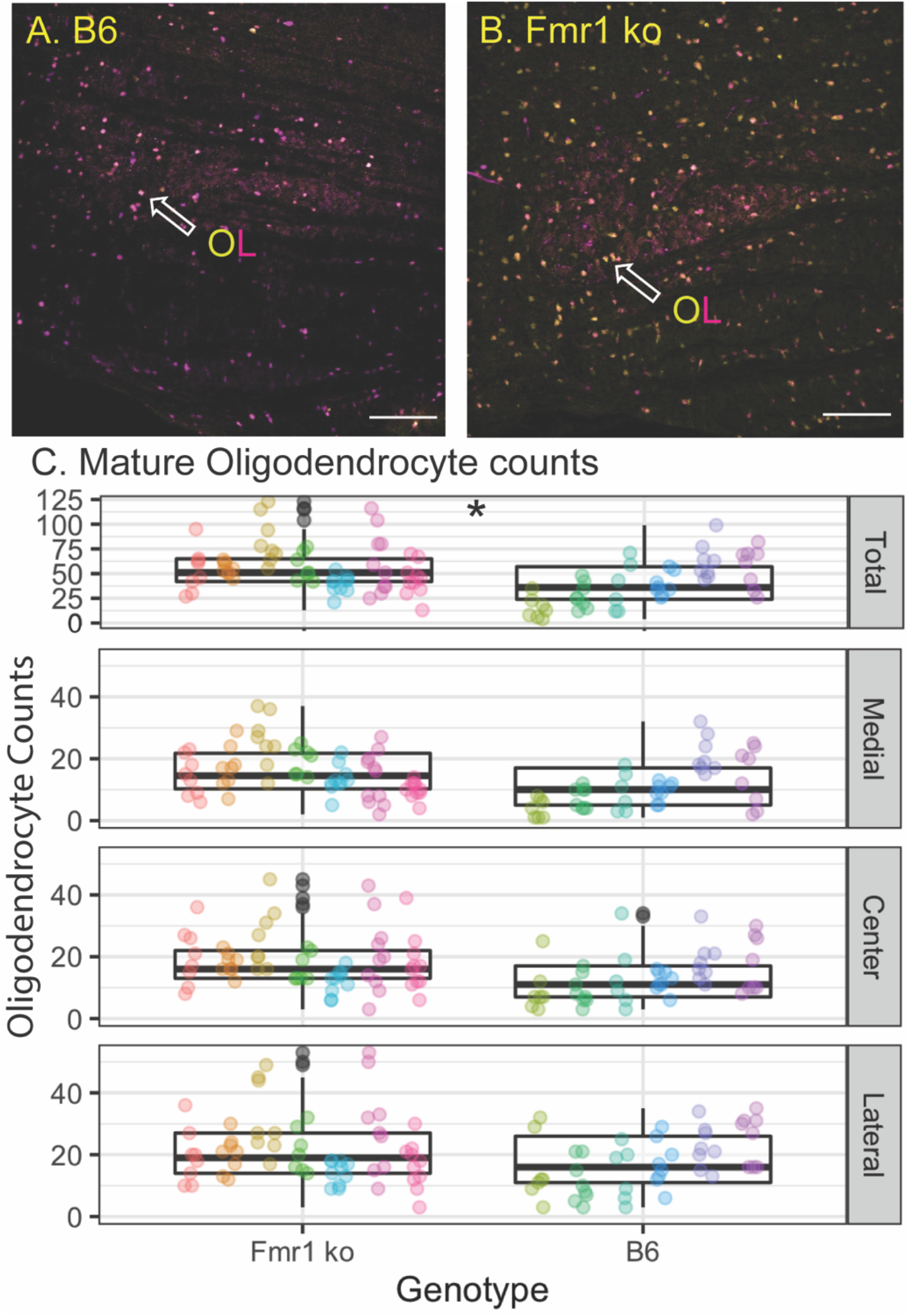
Oligodendrocyte number is increased in the MNTB independent of tonotopy in Fmr1 KO mice compared to B6. Representative images for OLs (yellow/magenta ASPA/sox-10 marker, arrowhead) in B6 (A) and Fmr1 KO (B) mice. Scale bar is 100 µm. Box plots show data from each individual measurement for Fmr1 KO (left) and B6 (right) for each animal (as indicated by different colors within the boxplots) for the total, medial, center, and lateral MNTB (C). N is 7 Fmr1 KO animals and 6 B6 animals. * = *p* < 0.05.

**Figure 6.**
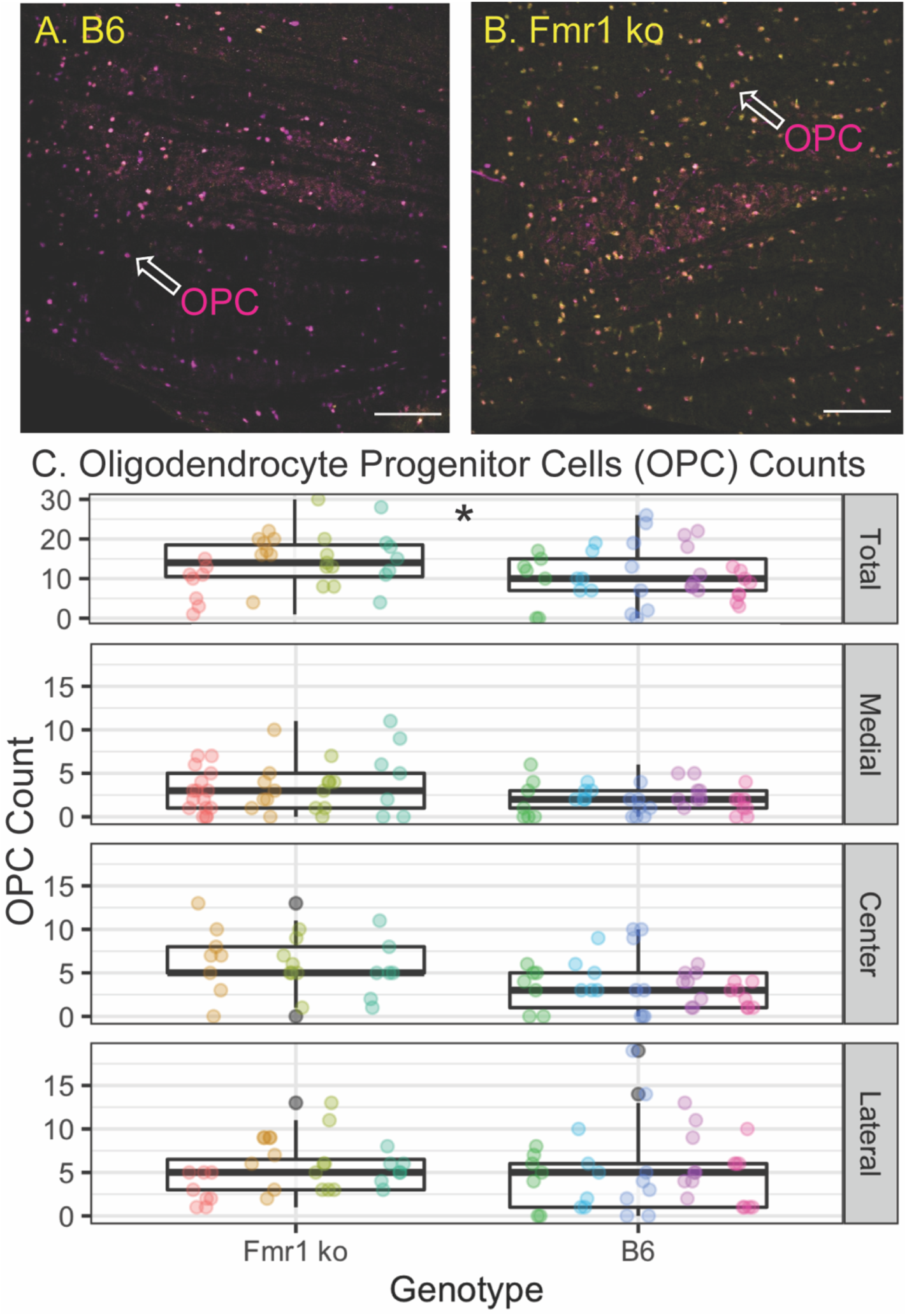
OPC number is increased in the MNTB independent of tonotopy in Fmr1 KO mice compared to B6. Representative images for OPCs (magenta sox-10 marker, arrowhead without ASPA (yellow) staining) in B6 and Fmr1 KO (B) mice. Scale bar is 100 µm. Box plots show data from each individual measurement for Fmr1 KO (left) and B6 (right) for each animal (as indicated by different colors within the boxplots) for the total, medial, center, and lateral MNTB (C). N is 4 Fmr1 KO animals and 5 B6 animals. * = *p* < 0.05.

### 4.4 Number of precursor oligodendrocytes

Other brain areas (cerebellum, neocortex, and others) show changes in myelination in FXS related to impaired maturation or function of oligodendrocyte precursor cells (Wang et al., 2005; Pacey et al., 2013; Jeon et al., 2017). Like OL count measures, the MNTB was divided into medial, central, and lateral subsections (Figure 6C), with representative images of the entire MNTB shown in Figure 6 for B6 (6A) and Fmr1 KO (6B). As described above, OPCs were quantified as cells that were positive for the sox-10 (magenta) marker without ASPA (yellow) staining (Figure 6A, B arrowheads). Like OL counts, there was no significant difference between Fmr1 KO and B6 mice based on tonotopic location (medial, center, or lateral, Figure 6C), however when data was summed for a total MNTB count, there was a significant increase in OPC number in Fmr1 KO mice compared to B6 (Figure 6C, top panel). Consistent with data from OLs, whole nucleus Q counts were consistent with raw counts such that there was no difference between genotypes based on tonotopy, but total estimates were significantly higher in Fmr1 KO animals compared to B6 (*p* < 0.0014). Together, these OPC and OL data show that changes in myelination observed through EM and CARS are not due to inherent decreases in numbers of mature or immature oligodendrocytes, however there are changes in overall number of these cells compared to wildtype animals.

## 5 Discussion

Mechanisms underlying auditory symptoms among patients with FXS are still poorly understood. The auditory brainstem is the first location along the ascending auditory pathway where sound information is processed, and it is likely key to identifying auditory difficulties since any changes at this level alter midbrain and cortical processing as well. Here we show decreased myelination in adult FXS mice (fiber diameter, axon diameter, thickness, and increased g-ratio) compared to wildtype in the region of the MNTB, which contains one of the largest and most heavily myelinated axons in the central nervous system. One possible mechanism for impaired myelination is changes in oligodendrocyte number and type. We show an increase in mature and precursor oligodendrocytes, suggesting perhaps an overcompensation for earlier reductions in OL numbers that others have shown (Pacey et al., 2013). Results from these measures suggest impaired myelination in the MNTB of Fmr1 KO mice, which could be a potential mechanism for auditory brainstem physiological and behavioral phenotypes.

While this is the first study to examine myelination changes in the auditory brainstem in FXS, previous work has shown reduced myelin sheath growth, decreases in total myelin (Doll et al., 2020), fewer myelinated and thinner axons, reductions in myelin basic protein, and development impacts of loss of FMRP (Pacey et al., 2013). FMRP is present in both precursor and mature oligodendrocytes (Wang et al., 2004; Giampetruzzi et al., 2013) and located subcellularly within myelin sheaths (Doll et al., 2020). However, recent work suggests that FMRP is not acting directly to regulate myelin basic protein (MBP) or other myelin protein transcripts, suggesting a diversity of roles for FMRP in myelin and oligodendrocyte dynamics (Giampetruzzi et al., 2013; Doll et al., 2020). Interestingly, inconsistent with the literature, and unexpectedly based on reductions in myelin characteristics measured here, we saw an increase in number of both OLs and OPCs. These differences can potentially be explained by an overcompensation for OL number reductions during development in this brain area, or an increased number of OLs that are not necessarily myelinating efficiently (fewer myelin sheaths per OL, non-myelinating OLs, etc.). In addition, it is possible that there are changes in proliferation ability of OPCs and OLs that were not measured here with only single time point measurements (Bu et al., 2004). What consequences these reductions in myelination have on conduction speed and physiological properties of the Calyx of Held in synaptic transmission are potential areas for future research.

G-ratio is a common measure for not just structural but potential physiological properties of myelin conduction along an axon. The g-ratios we measured for Fmr1 KO and B6 animals are similar to previous work (particularly for wildtype B6 animals) in the MNTB (Ford et al., 2015; Stange-Marten et al., 2017). Conduction velocity is an important factor in reliable processing of sound location information, resulting in the Calyx of Held which is highly specialized for speed and temporal fidelity. Increased g-ratio suggests slower conduction velocities and thinner myelin in Fmr1 KO mice (Rushton, 1951; Berman et al., 2019). However note that there may be other factors which contribute to conduction velocity, for example Ranvier node and internode spacing, length, and diameter (Ford et al., 2015). In addition, while we did not see differences in myelin properties based on location or tonotopic area, it is possible that node dynamics vary more with tonotopic area while myelin thickness and diameter are less impacted based on tonotopy. However, note that such differences may also be explained by the relative importance of ILD versus ITD processing in different animal models. The previous work showing tonotopic differences was performed in gerbils, a rodent model suitable to study both low and high-frequency hearing. In that animal model, one important role for MNTB neurons is to provide inhibitory input to ITD processing (Ford et al., 2015). By contrast, mice are primarily high-frequency specialists. Consistent with their hearing spectrum and the fact that mice primarily use ILD cues, previous work has shown that there are no differences in myelination properties based on tonotopy (Stange-Marten et al., 2017), similar to what we see in this study.

In the current study we did not directly label Calyx of Held axons. Rather, we made measurements within the MNTB, measuring axons that were projecting in the coronal plane. Although axons projecting to MNTB are significantly larger than any other passing fibers and can be discerned by eye relatively easily and reliably, we cannot rule out the possibility that some measured axons may be “passing through”. If our analysis did include some passing fibers, these would likely decrease the observed differences between wild type and mutant, such that the results presented in this study may be a lower limit of the true differences between wild type and mutant mouse model. This would also limit the interpretation of conclusions about tonotopic location of the measurements and whether indeed there are differences based on frequency-coding. Note that CARS nor EM techniques inherently discriminate between different types of axons. However, based on the size of the axons that we measured in both EM and CARS, fibers are consistent with the expected size of Calyx axons in the same area (Ford et al., 2015; Sinclair, 2017; Stange-Marten et al., 2017), therefore we are confident that at least the large majority of the axons are Calyceal projections.

This is the first study to show myelination changes in the auditory brainstem sound localization pathway in FXS mice. Thinner and smaller diameter Calyx axons with increased g-ratio may underly sound localization and auditory hypersensitivity issues seen behaviorally in mice and described by patient’s with FXS. Interestingly, we saw an increase in both mature and immature OLs suggesting that myelin deficits are not due simply to fewer myelinating OLs. Future studies elucidating when during development myelin deficits begin and how OLs develop from precursors into mature OLs in this brain area would be crucial to understanding the complete picture of auditory brainstem phenotypes in FXS.

## 7 Author Contributions

All authors helped write and revise the manuscript. EAM, SP, AL, AK developed the ideas and methods. EAM, SP, and AL collected the data for the manuscript. EAM and AL created the statistical analyses and figures for the manuscript.

## 9 Acknowledgements

We thank Dr. Jennifer Bourne for EM imaging. We also gratefully thank Clara Bacmeister and Ethan Hughes (University of Colorado, Anschutz) for providing antibodies, staining protocols, and overall support of this project.

## References

Barak, B., Zhang, Z., Liu, Y., Nir, A., Trangle, S. S., Ennis, M., et al. (2019). Neuronal deletion of Gtf2i, associated with Williams syndrome, causes behavioral and myelin alterations rescuable by a remyelinating drug. Nat Neurosci 22, 700–708. doi:10.1038/s41593-019-0380-9.

Bates, D., Mächler, M., Bolker, B., and Walker, S. (2014). Fitting Linear Mixed-Effects Models using lme4. arXiv.

Bear, M. F., Huber, K. M., and Warren, S. T. (2004). The mGluR theory of fragile X mental retardation. Trends in Neurosciences 27, 370–377. doi:10.1016/j.tins.2004.04.009.

Berman, S., Filo, S., and Mezer, A. A. (2019). Modeling conduction delays in the corpus callosum using MRI-measured g-ratio. Neuroimage 195, 128–139. doi:10.1016/j.neuroimage.2019.03.025.

Bitto, E., Bingman, C. A., Wesenberg, G. E., McCoy, J. G., and Phillips, G. N. (2007). Structure of aspartoacylase, the brain enzyme impaired in Canavan disease. Proceedings of the National Academy of Sciences 104, 456–461. doi:10.1073/pnas.0607817104.

Brown, M. R., Kronengold, J., Gazula, V.-R., Chen, Y., Strumbos, J. G., Sigworth, F. J., et al. (2010). Fragile X mental retardation protein controls gating of the sodium-activated potassium channel Slack. Nature Neuroscience 13, 819–821. doi:10.1038/nn.2563.

Bu, J., Banki, A., Wu, Q., and Nishiyama, A. (2004). Increased NG2+ glial cell proliferation and oligodendrocyte generation in the hypomyelinating mutant shiverer. Glia 48, 51–63. doi:10.1002/glia.20055.

Chomiak, T., and Hu, B. (2009). What Is the Optimal Value of the g-Ratio for Myelinated Fibers in the Rat CNS? A Theoretical Approach. PLOS ONE 4, e7754. doi:10.1371/journal.pone.0007754.

Curry, R. J., Peng, K., and Lu, Y. (2018). Neurotransmitter- and Release-Mode-Specific Modulation of Inhibitory Transmission by Group I Metabotropic Glutamate Receptors in Central Auditory Neurons of the Mouse. The Journal of Neuroscience 38, 8187–8199. doi:10.1523/JNEUROSCI.0603-18.2018.

Dahlhaus, R. (2018). Of Men and Mice: Modeling the Fragile X Syndrome. Front Mol Neurosci 11. doi:10.3389/fnmol.2018.00041.

Doll, C. A., Yergert, K. M., and Appel, B. H. (2020). The RNA binding protein fragile X mental retardation protein promotes myelin sheath growth. Glia 68, 495–508. doi:10.1002/glia.23731.

El-Hassar, L., Song, L., Tan, W. J. T., Large, C. H., Alvaro, G., Santos-Sacchi, J., et al. (2019). Modulators of Kv3 Potassium Channels Rescue the Auditory Function of Fragile X Mice. The Journal of Neuroscience 39, 4797–4813. doi:10.1523/JNEUROSCI.0839-18.2019.

Ford, M. C., Alexandrova, O., Cossell, L., Stange-Marten, A., Sinclair, J., Kopp-Scheinpflug, C., et al. (2015). Tuning of Ranvier node and internode properties in myelinated axons to adjust action potential timing. Nat Commun 6, 8073. doi:10.1038/ncomms9073.

Gantois, I., Khoutorsky, A., Popic, J., Aguilar-Valles, A., Freemantle, E., Cao, R., et al. (2017). Metformin ameliorates core deficits in a mouse model of fragile X syndrome. Nat. Med. 23, 674–677. doi:10.1038/nm.4335.

Garcia-Pino, E., Gessele, N., and Koch, U. (2017). Enhanced Excitatory Connectivity and Disturbed Sound Processing in the Auditory Brainstem of Fragile X Mice. J. Neurosci. 37, 7403–7419. doi:10.1523/JNEUROSCI.2310-16.2017.

Giampetruzzi, A., Carson, J. H., and Barbarese, E. (2013). FMRP and myelin protein expression in oligodendrocytes. Mol. Cell. Neurosci. 56, 333–341. doi:10.1016/j.mcn.2013.07.009.

Grothe, B., Pecka, M., and McAlpine, D. (2010). Mechanisms of sound localization in mammals. Physiol. Rev. 90, 983–1012. doi:10.1152/physrev.00026.2009.

Gundersen, H. J., Jensen, E. B., Kiêu, K., and Nielsen J, null (1999). The efficiency of systematic sampling in stereology--reconsidered. J Microsc 193, 199–211. doi:10.1046/j.1365-2818.1999.00457.x.

Hagerman, P. J., and Hagerman, P. J. (2008). The fragile X prevalence paradox. J. Med. Genet. 45, 498–499. doi:10.1136/jmg.2008.059055.

Hagerman, R. J., Berry-Kravis, E., Kaufmann, W. E., Ono, M. Y., Tartaglia, N., Lachiewicz, A., et al. (2009). Advances in the treatment of fragile X syndrome. Pediatrics 123, 378–390. doi:10.1542/peds.2008-0317.

Imamura, O., Arai, M., Dateki, M., Oishi, K., and Takishima, K. (2020). Donepezil-induced oligodendrocyte differentiation is mediated through estrogen receptors. J Neurochem 155, 494–507. doi:10.1111/jnc.14927.

Jeon, S. J., Ryu, J. H., and Bahn, G. H. (2017). Altered Translational Control of Fragile X Mental Retardation Protein on Myelin Proteins in Neuropsychiatric Disorders. Biomol Ther (Seoul) 25, 231–238. doi:10.4062/biomolther.2016.042.

Kim, H., Gibboni, R., Kirkhart, C., and Bao, S. (2013a). Impaired Critical Period Plasticity in Primary Auditory Cortex of Fragile X Model Mice. J Neurosci 33, 15686–15692. doi:10.1523/JNEUROSCI.3246-12.2013.

Kim, J. H., Renden, R., and von Gersdorff, H. (2013b). Dysmyelination of Auditory Afferent Axons Increases the Jitter of Action Potential Timing during High-Frequency Firing. Journal of Neuroscience 33, 9402–9407. doi:10.1523/JNEUROSCI.3389-12.2013.

Kulesza Jr., R. J., Lukose, R., and Stevens, L. V. (2011). Malformation of the human superior olive in autistic spectrum disorders. Brain Research 1367, 360–371. doi:10.1016/j.brainres.2010.10.015.

Kulesza, R. J., and Mangunay, K. (2008). Morphological features of the medial superior olive in autism. Brain Research 1200, 132–137. doi:10.1016/j.brainres.2008.01.009.

Larson, V. A., Mironova, Y., Vanderpool, K. G., Waisman, A., Rash, J. E., Agarwal, A., et al. (2018). Oligodendrocytes control potassium accumulation in white matter and seizure susceptibility. eLife 7, e34829. doi:10.7554/eLife.34829.

Lenth, R. (2019). emmeans: Estimated Marginal Means, aka Least-Squares Means. Available at: https://CRAN.R-project.org/package=emmeans.

Lu, Y. (2019). Subtle differences in synaptic transmission in medial nucleus of trapezoid body neurons between wild-type and Fmr1 knockout mice. Brain Research 1717, 95–103. doi:10.1016/j.brainres.2019.04.006.

McCullagh, E. A., Poleg, S., Greene, N. T., Huntsman, M. M., Tollin, D. J., and Klug, A. (2020a). Characterization of Auditory and Binaural Spatial Hearing in a Fragile X Syndrome Mouse Model. eNeuro 7. doi:10.1523/ENEURO.0300-19.2019.

McCullagh, E. A., Rotschafer, S. E., Auerbach, B. D., Klug, A., Kaczmarek, L. K., Cramer, K. S., et al. (2020b). Mechanisms underlying auditory processing deficits in Fragile X syndrome. The FASEB Journal 00, 1–18. doi:https://doi.org/10.1096/fj.201902435R.

McCullagh, E. A., Salcedo, E., Huntsman, M. M., and Klug, A. (2017). Tonotopic alterations in inhibitory input to the medial nucleus of the trapezoid body in a mouse model of Fragile X syndrome. Journal of Comparative Neurology 262, 375. doi:10.1002/cne.24290.

Orthmann-Murphy, J., Call, C. L., Molina-Castro, G. C., Hsieh, Y. C., Rasband, M. N., Calabresi, P. A., et al. (2020). Remyelination alters the pattern of myelin in the cerebral cortex. eLife 9, e56621. doi:10.7554/eLife.56621.

Osterweil, E. K., Chuang, S.-C., Chubykin, A. A., Sidorov, M., Bianchi, R., Wong, R. K. S., et al. (2013). Lovastatin corrects excess protein synthesis and prevents epileptogenesis in a mouse model of fragile X syndrome. Neuron 77, 243–250. doi:10.1016/j.neuron.2012.01.034.

Pacey, L. K. K., Xuan, I. C. Y., Guan, S., Sussman, D., Henkelman, R. M., Chen, Y., et al. (2013). Delayed myelination in a mouse model of fragile X syndrome. Hum. Mol. Genet. 22, 3920–3930. doi:10.1093/hmg/ddt246.

Phan, B. N., Bohlen, J. F., Davis, B. A., Ye, Z., Chen, H.-Y., Mayfield, B., et al. (2020). A myelin-related transcriptomic profile is shared by Pitt–Hopkins syndrome models and human autism spectrum disorder. Nature Neuroscience 23, 375–385. doi:10.1038/s41593-019-0578-x.

R Core Team (2013). R: A Language and Environment for Statistical Computing. Vienna, Austria: R Foundation for Statistical Computing Available at: http://www.R-project.org/.

Rajaratnam, A., Shergill, J., Salcedo-Arellano, M., Saldarriaga, W., Duan, X., and Hagerman, R. (2017). Fragile X syndrome and fragile X-associated disorders. F1000Research 6, 2112. doi:10.12688/f1000research.11885.1.

Rotschafer, S. E., and Cramer, K. S. (2017). Developmental Emergence of Phenotypes in the Auditory Brainstem Nuclei of Fmr1Knockout Mice. eNeuro 4, 1–21. doi:10.1523/ENEURO.0264-17.2017.

Rotschafer, S. E., Marshak, S., and Cramer, K. S. (2015). Deletion of Fmr1 Alters Function and Synaptic Inputs in the Auditory Brainstem. PLoS ONE 10, 1–15. doi:10.1371/journal.pone.0117266.

Rushton, W. A. H. (1951). A theory of the effects of fibre size in medullated nerve. J Physiol 115, 101–122.

Schindelin, J., Arganda-Carreras, I., Frise, E., Kaynig, V., Longair, M., Pietzsch, T., et al. (2012). Fiji: an open-source platform for biological-image analysis. Nature Methods 9, 676–682. doi:10.1038/nmeth.2019.

Schmitz, C., and Hof, P. R. (2005). Design-based stereology in neuroscience. Neuroscience 130, 813–831. doi:10.1016/j.neuroscience.2004.08.050.

Sinclair, J. L. (2017). Sound-Evoked Activity Influences Myelination of Brainstem Axons in the Trapezoid Body. Journal of Neuroscience 37, 8239–8255. doi:10.1523/JNEUROSCI.3728-16.2017.

Stange-Marten, A., Nabel, A. L., Sinclair, J. L., Fischl, M., Alexandrova, O., Wohlfrom, H., et al. (2017). Input timing for spatial processing is precisely tuned via constant synaptic delays and myelination patterns in the auditory brainstem. Proc Natl Acad Sci U S A 114, E4851–E4858. doi:10.1073/pnas.1702290114.

Strumbos, J. G., Brown, M. R., Kronengold, J., Polley, D. B., and Kaczmarek, L. K. (2010). Fragile X Mental Retardation Protein Is Required for Rapid Experience-Dependent Regulation of the Potassium Channel Kv3.1b. Journal of Neuroscience 30, 10263–10271. doi:10.1523/JNEUROSCI.1125-10.2010.

The Dutch-Belgian Fragile X Consorthium, Bakker, C. E., Verheij, C., Willemsen, R., van der Helm, R., Oerlemans, F., et al. (1994). Fmr1 knockout mice: A model to study fragile X mental retardation. Cell 78, 23–33. doi:10.1016/0092-8674(94)90569-X.

Tian, Y., Yang, C., Shang, S., Cai, Y., Deng, X., Zhang, J., et al. (2017). Loss of FMRP Impaired Hippocampal Long-Term Plasticity and Spatial Learning in Rats. Front Mol Neurosci 10, 269. doi:10.3389/fnmol.2017.00269.

Till, S. M., Asiminas, A., Jackson, A. D., Katsanevaki, D., Barnes, S. A., Osterweil, E. K., et al. (2015). Conserved hippocampal cellular pathophysiology but distinct behavioural deficits in a new rat model of FXS. Hum Mol Genet 24, 5977–5984. doi:10.1093/hmg/ddv299.

Wang, H., Fu, Y., Zickmund, P., Shi, R., and Cheng, J.-X. (2005). Coherent Anti-Stokes Raman Scattering Imaging of Axonal Myelin in Live Spinal Tissues. Biophysical Journal 89, 581–591. doi:10.1529/biophysj.105.061911.

Wang, H., Ku, L., Osterhout, D. J., Li, W., Ahmadian, A., Liang, Z., et al. (2004). Developmentally-programmed FMRP expression in oligodendrocytes: a potential role of FMRP in regulating translation in oligodendroglia progenitors. Hum. Mol. Genet. 13, 79–89. doi:10.1093/hmg/ddh009.

Wang, Y., Sakano, H., Beebe, K., Brown, M. R., de Laat, R., Bothwell, M., et al. (2014). Intense and specialized dendritic localization of the fragile X mental retardation protein in binaural brainstem neurons: A comparative study in the alligator, chicken, gerbil, and human: FMRP localization in NL/MSO dendrites. Journal of Comparative Neurology 522, 2107–2128. doi:10.1002/cne.23520.

Weatherstone, J. H., Kopp-Scheinpflug, C., Pilati, N., Wang, Y., Forsythe, I. D., Rubel, E. W., et al. (2016). Maintenance of neuronal size gradient in MNTB requires sound-evoked activity. Journal of Neurophysiology 117, 756–766. doi:10.1152/jn.00528.2016.

Wickham, H. (2016). ggplot2: Elegant Graphics for Data Analysis. Springer-Verlag New York Available at: http://ggplot2.org.

Zorio, D. A. R., Jackson, C. M., Liu, Y., Rubel, E. W., and Wang, Y. (2017). Cellular distribution of the fragile X mental retardation protein in the mouse brain: Fmrp distribution in the mouse brain. Journal of Comparative Neurology 525, 818–849. doi:10.1002/cne.24100.

Zuo, H., Wood, W. M., Sherafat, A., Hill, R. A., Lu, Q. R., and Nishiyama, A. (2018). Age-Dependent Decline in Fate Switch from NG2 Cells to Astrocytes After Olig2 Deletion. J Neurosci 38, 2359–2371. doi:10.1523/JNEUROSCI.0712-17.2018.

